# Defining the single base importance of human mRNAs and lncRNAs

**DOI:** 10.1101/2023.06.12.544536

**Authors:** Rui Fan, Xiangwen Ji, Jianwei Li, Qinghua Cui, Chunmei Cui

**Affiliations:** Department of Physiology and Pathophysiology, Department of Biomedical Informatics, Center for Noncoding RNA Medicine, MOE Key Lab of Cardiovascular Sciences, School of Basic Medical Sciences, Peking University, 38 Xueyuan Rd, Beijing, 100191, China; Institute of Computational Medicine, School of Artificial Intelligence, Hebei University of Technology, Tianjin, 300401, China

**Keywords:** single base importance, algorithm, tumor mutation burden, SARS-CoV-2

## Abstract

As the fundamental unit of a gene and its transcripts, nucleotides have enormous impacts on molecular function and evolution, and thus on phenotypes and diseases. Given that different nucleotides on one gene often exhibit diverse levels of effects, it is quite crucial to comprehensively and quantitatively measure the importance of each base on a gene transcript, however, tools are still not available. Here we proposed Base Importance Calculator (BIC), an algorithm to calculate the importance score of single bases based on sequence information of human mRNAs and long noncoding RNAs (lncRNAs). We then confirmed its power by applying BIC to three different tasks. Firstly, we revealed that BIC can effectively evaluate the pathogenicity of both genes and single bases by analyzing the BIC scores and the pathogenicity of single nucleotide variations (SNVs). Moreover, the BIC score in the Cancer Genome Atlas (TCGA) somatic mutations is able to predict the prognosis of some cancers. Finally, we show that BIC can also precisely predict the transmissibility of SARS-CoV-2. The above results indicate that BIC is a useful tool for evaluating the single base important of human mRNAs and lncRNAs.

**Key Points:**

- BIC could measure the single base importance of human mRNAs and lncRNAs.
- BIC could be applied to many aspects including measuring the pathogenicity of SNVs and enhancing the ability of predicting cancer survival.
- BIC could predict the transmissibility of SARS-CoV-2

## INTRODUCTION

Nucleotides represent the basic block of the sequence of DNA or RNA. Given the central role of the sequence-structure-function paradigm, it is no surprise that nucleotides have enormous impacts on molecular function and evolution, and thus on phenotypes and diseases. Nucleotides located at different position of one gene always play distinct roles, such as the splice conjunction sites, the nucleotides on 3’UTR deciding the binding of miRNA^1^, and the nucleotides on a highly conserved region across species usually involving in substantial biological processes including DNA replication^2^, transcription^3^, translation^4^, and regulation of gene expression^5^. It is thus reasonable to assume that each individual nucleotide of a gene would make different contributions, and identification of these key nucleotides could facilitate locating the biologically functional sites of a gene ^6,7^ and even developing new therapeutic target^8^.

At present, genome-wide association studies (GWASs) assists in identifying the mutations associated with a risk for diseases or traits^9^. And omics techniques such as whole genome sequencing (WGS), whole exome sequencing (WES), and RNA-sequencing have detected actual single nucleotide variations (SNVs)^10^, the most abundant and simplest type of germline variations and somatic mutations^11,12^. However, due to the factors such as sample size and SNV density, it is still challenging to recognize all significant SNVs using above technics. Therefore, it is imperative to devise a computational method to comprehensively and quantitatively evaluate the importance of each base on a gene for prioritizing those significant sites.

There are growing algorithms available for accessing the functional impacts of multiple types of SNVs, for example, SIFT^13^ is specifically used for nonsynonymous SNVs, SilVA^14^ is devised for synonymous variants, and FATHMM-MKL^15^ is a more general predictors applied to all types of SNVs. Despite that current tools are convenient in evaluating all SNVs, with the merit of gene length to the number of SNVs, a large proportion of sites on a gene remain unquantified for their functional effects. In addition, above methods are not applicable on measuring the importance of actual variants or single base on lncRNAs, which are also demonstrated to play important roles in a variety of diseases^16,17^.

Generally, there are no efficient computational algorithms available to comprehensively and quantitatively evaluate single base importance for all bases of a mRNA and/or lncRNA sequence and it is thus emergently necessary to develop a tool for the above purpose. In this paper, we designed an algorithm, Base Importance Calculator (BIC), based on the GIC (Gene Importance Calculator) algorithm^18^ which is developed to efficiently measure the essentiality of both mRNA and lncRNA sequences. The BIC score of one given base is calculated by evaluating the average change of the GIC scores when mutating this base to the other three bases. Subsequently, the effectiveness of the BIC algorithm and tool was confirmed by three different tasks. Firstly, we showed that the BIC score is able to efficiently predict the pathogenicity of both mRNAs and lncRNAs. Then, we found that the BIC score can precisely predict the cancer prognosis by weighting the Tumor Mutation Burden (TMB). Finally, BIC showed a powerful ability to predict the transmissibility of different SARS-CoV-2 strains. All the results show that BIC is a useful metric for evaluating the single base important of human mRNAs and lncRNAs.

## METHODS AND MATERIALS

### Data collection

To confirm whether the BIC score can predict pathogenicity or not, we downloaded the disease-related SNVs from four databases including ClinVar^19^, the Human Gene Mutation Database (HGMD)^20^, the Catalogue of Somatic Mutations in Cancer (COSMIC)^21^, and the Cancer Genome Atlas (TCGA)^22^. We downloaded these databases in VCF or MAF format, which contains the SNV locus and its relevant information. As some datasets were annotated using different genome assemblies, we employed the liftovervcf tool from the Genome Analysis Toolkit (GATK) package^23^ to convert all datasets to the GRCh38 genome assembly, which was obtained from UCSC Genome Browser. We obtained human lncRNA transcripts from the NONCODE database^24^. Overall, we collected 1,338,130 records from ClinVar, 111,152 records from HGMD, 15,081,012 records from COSMIC, and 3,276,147 records from TCGA. In addition, we downloaded other metrics, including phastCons from UCSC Genome Browser and dNdS with the mouse from Ensembl, for comparison with other metrics. Furthermore, since BIC relies on GIC algorithm, we downloaded a standalone version of GIC tool from the website (https://www.cuilab.cn/gic/download). For SARS-CoV-2, we obtained the lineage definitions and doubling times from NCBI Virus^25^.

### Calculating single base importance

In this study, we evaluated the importance of a given single base by calculating the average difference of GIC score between the wild type sequence and the mutant sequences of this base. The process of calculating the BIC score is describe in the following formula:

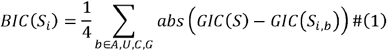

where S refers to the wild type sequence, S_i_ represents the i^th^ base in sequence S, S_i,b_ denotes the mutant sequence obtained by changing the i^th^ base to b, and GIC is the function used to calculate the importance score of the sequence S. A base with a higher BIC score indicates that changing it to another nucleotide will cause a greater change in the corresponding GIC score, which could disrupt the function of the gene more seriously.

As shown in Figure 1, specifically, to calculate the importance score of a given single base (the blue ‘A’), we generate a set of variant sequences by altering that base with the other three bases (‘T’, ‘C’, and ‘G’), and then calculate the absolute mean deviation between the wild type sequence and each mutant sequence. By this way, we can calculate the single base importance score for each base of the gene transcript sequence.

**Figure 1.**
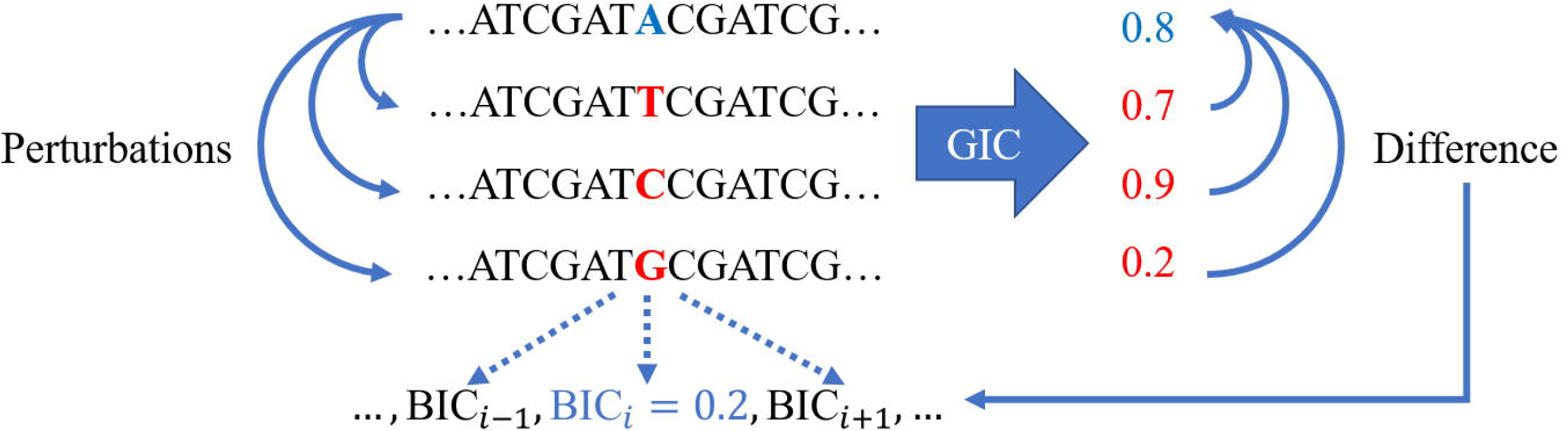
Diagram of defining the BIC score for a given single base

### Defining gene-level importance score with base importance scores

As we utilized the absolute value of the relative change between the GIC scores of the wild type sequence and the mutant sequences, the BIC score does not follow a normal distribution. Therefore, to define the gene-level BIC score (gBIC), we used the median value of all BIC scores for a given gene transcript sequence, which is described as follows:

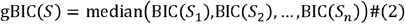

where n is the length of the sequence.

### Defining BIC derived Tumor Mutation Burden (TMB)

The Tumor Mutation Burden (TMB) was defined as:

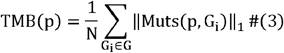

where p refers to a single patient, G represents all the measured genes, Muts is a function that returns all mutated sequences in gene i of patient p, and N is a normalization factor representing the total length of all genes here. TMB is simply the count of mutations per unit sequence length. To evaluate the efficiency of BIC measurement, we introduced three versions of BIC derived TMB, defined by the following equation:

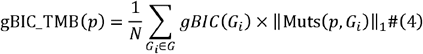

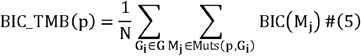

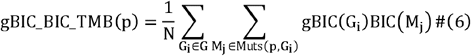

The first two scores are weighted by its gBIC or BIC score and the last one is the combination of first two.

### Statistical methods

Due to the non-normal distribution of both BIC and gBIC scores, non-parametric statistical methods were used for base-level and gene-level comparison. Specifically, Spearman’s coefficient was utilized for correlation analysis, while the Mann-Whitney U rank test and the Wilcoxon signed-rank test were employed for unpaired and paired comparisons between two groups. Additionally, to analyze survival rates, the Kaplan-Meier curve was utilized, and both the Cox regression log likelihood ratio test and Log-rank test were used to assess the significance of the results. These statistical methods were chosen for their ability to provide robust and accurate analysis in the presence of non-normal data distributions.

## RESULTS

### BIC is able to predict the pathogenicity of a gene

To confirm the accuracy and effectiveness of the BIC algorithm, we calculated the gBIC scores of genes from human pathological gene mutation databases, including ClinVar, HGMD, COSMIC, and TCGA. We then conducted a detailed analysis to explore the relationship between gBIC scores and the pathological mutation rate of coding genes. Here the pathological mutation rate of a gene is defined as the number of pathological mutations annotated in the database divided by the gene length. Specifically, for pathological mutation rate, we only considered single nucleotide substitution to ensure consistency in the calculation of the BIC score which currently does not take indel into account. As a result, we revealed a significant positive correlation between the gBIC score and the pathological mutation rate in all the four databases (p-value=3.49e-57 for ClinVar, p-value=7.17e-62 for HGMD, p-value<1.0e-323 for COSMIC, and p-value<1.0e-323 for TCGA; Figure 2, Supplementary Figure 1), suggesting that gBIC could be a strong indicator of the pathogenicity of coding genes.

**Figure 2.**
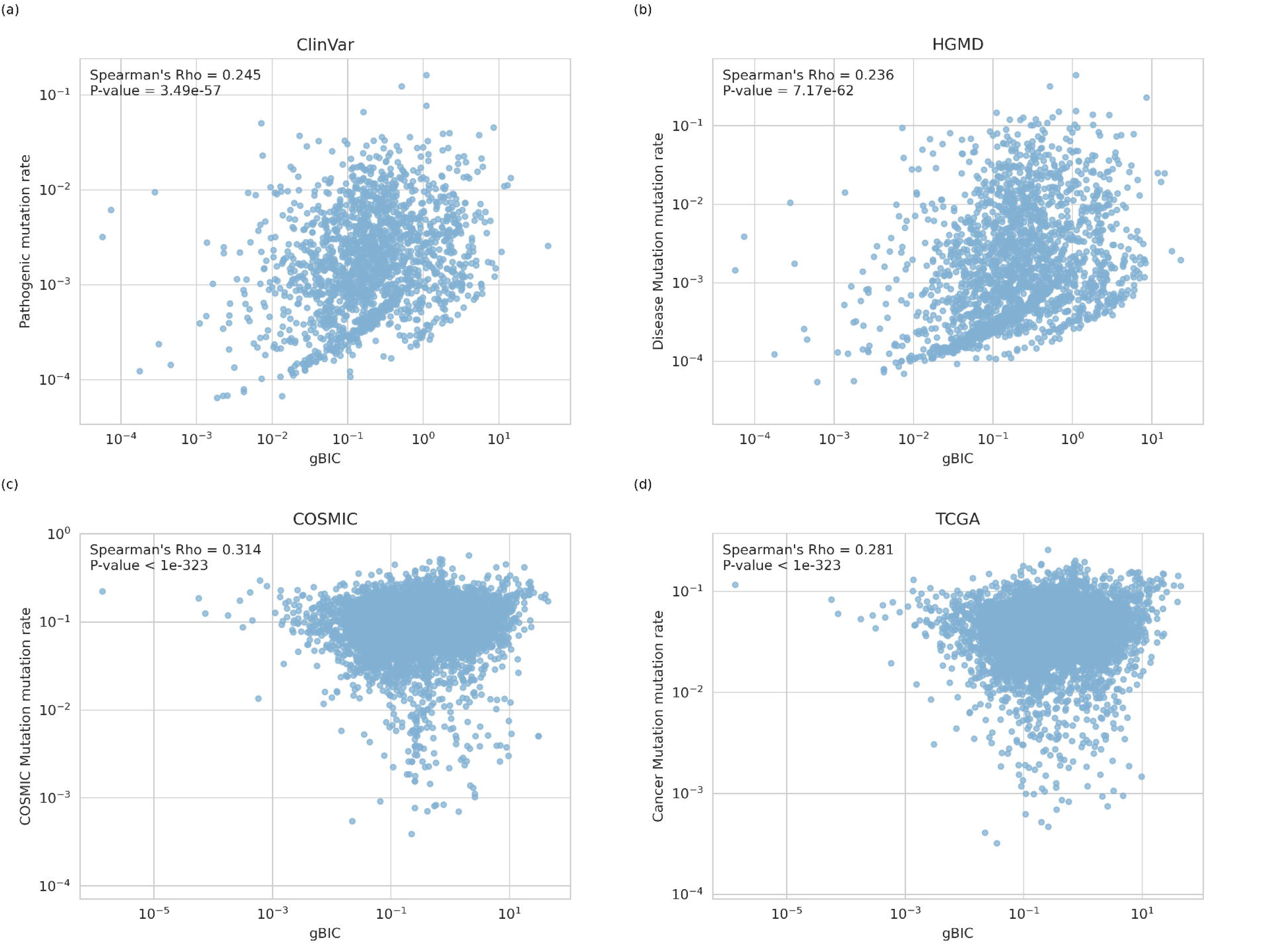
The correlation between gene-level BIC (gBIC) score and the pathological mutation rate for coding genes in databases of a) ClinVar, b) HGMD, c) COSMIC, and d) TCGA. The p-value was calculated by Spearman’s correlation analysis.

### BIC is able to predict pathological mutations in coding genes and long noncoding genes

The above results showed that the gene-level BIC score works effectively in predicting gene pathogenicity, we next ask whether the single base level BIC score can also measure the pathogenicity of single base. For doing so, we analyzed the BIC scores for the bases of the pathological mutation sites and their neighboring sites. As illustrated in Figure 3, we found that the bases hosting pathological mutations exhibit significantly higher BIC scores than their neighboring bases in all of the four databases (p-value<1.0e-323 and p-value=3.84e-168 for the 5’ and 3’ neighbor site of ClinVar, p-value<1.0e-323 and p-value=9.39e-156 for the 5’ and 3’ neighbor site of HGMD, p-value<1.0e-323 and p-value= p-value<1.0e-323 for the 5’ and 3’ neighbor site of both COSMIC and TCGA), suggesting that the BIC score may serve as a useful metric for assessing the pathogenicity of single base mutations.

**Figure 3.**
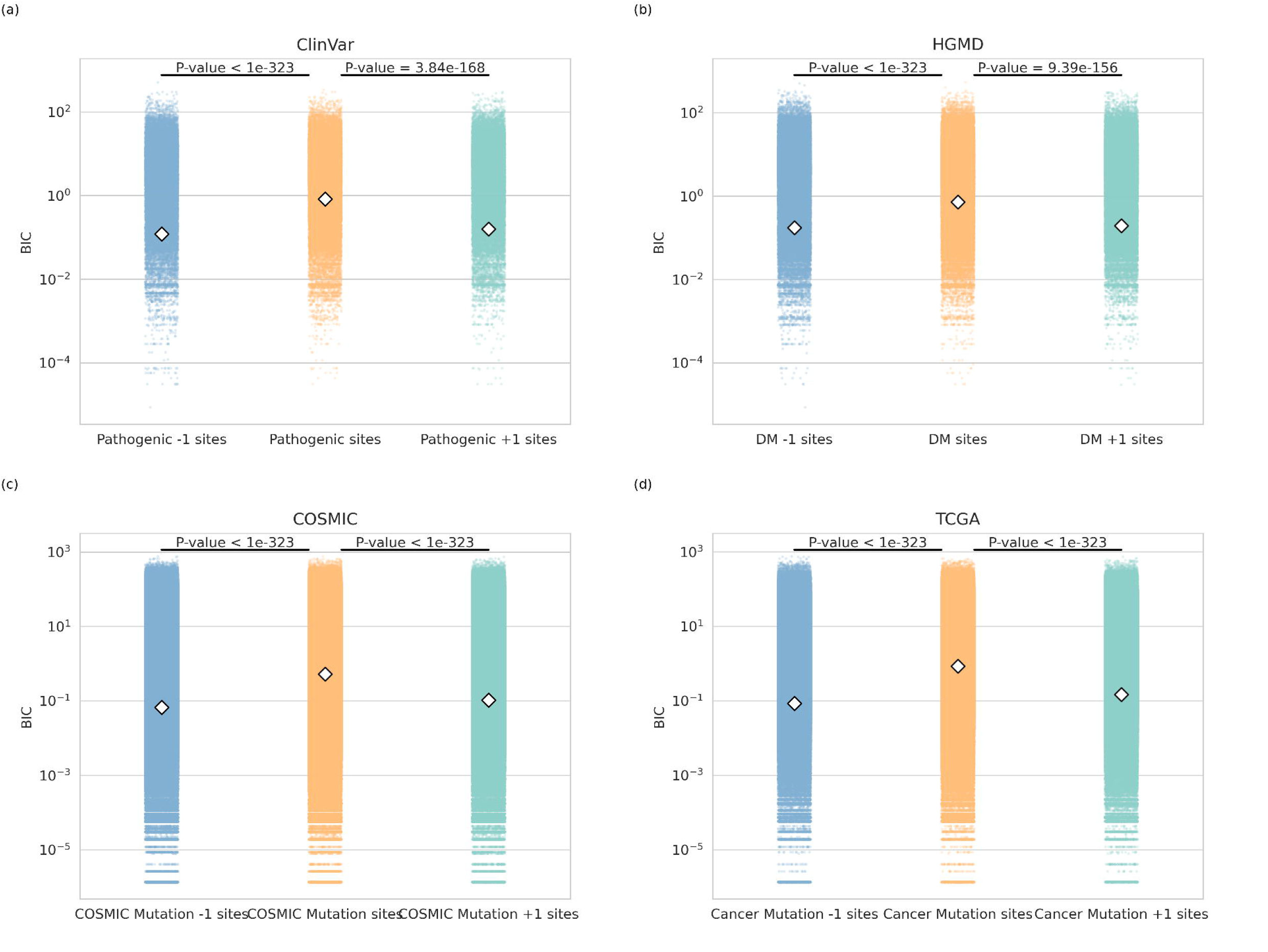
Comparison of the BIC scores of the bases hosting pathological mutation sites (yellow) and their upstream neighbor single bases (blue) and downstream neighbor single bases (green) for protein coding genes in databases of a) ClinVar, b) HGMD, c) COSMIC, and d) TCGA. The p-value was calculated by Mann-Whitney U rank test.

The above results showed that both the gBIC and BIC scores are significantly correlated with the mutational pathogenicity of protein coding genes. Moreover, it is well known noncoding RNAs make up a significant portion of the human genome, accounting for more than 90% of all genes, and research increasingly highlights their importance^26,27^. As BIC can be also used for analyzing lncRNA sequences, it is thus intriguing to investigate whether BIC algorithm still works well on lncRNAs. To address this question, we applied a similar analysis pipeline to human lncRNAs. As shown in Supplementary Figures 2-3, the results on lncRNAs are quite similar with those on the protein coding genes. BIC is also able to predict the pathogenicity of both lncRNAs and single bases in lncRNAs.

### BIC improves the effectiveness of Tumor Mutation Burden (TMB) in predicting the prognosis of some cancers

Tumor Mutation Burden (TMB) represents the mutation rate across all genes and is able to predict the response to immune checkpoint inhibitor treatment of certain cancers^28^. Patients with a higher TMB may have a greater likelihood of harboring key mutations that can be recognized and eliminated by the immune system. However, the limitation of TMB is that each mutation is treated equally, despite the fact that different mutations have varying effect. To address this, BIC derived TMB scores were defined to test whether these new TMB scores can improve the performance of survival prediction. The patients were then divided into two groups based on their median scores. The results indicate that in certain cancers, such as Brain Lower Grade Glioma (LGG, log-rank p-value=2.75e-8), Mesothelioma (MESO, log-rank p-value=1.2e-3) and Uveal Melanoma (UVM, log-rank p-value=5.47e-2), BIC-weighted TMB can significantly improve survival prediction efficiency (Figure 4 and Supplementary Figure 4). Additionally, gBIC improves the prognosis prediction of Rectum adenocarcinoma (READ, log-rank p-value=1.16e-2), while Bladder Urothelial Carcinoma (BLCA, log-rank p-value=1.43e-5) and Thymoma (THYM, log-rank p-value=6.74e-3) benefit from both gBIC and BIC scores in predicting survival time (Figure 4 and Supplementary Figure 4), suggesting that BIC is effective in improving the original TMB measurement.

**Figure 4.**
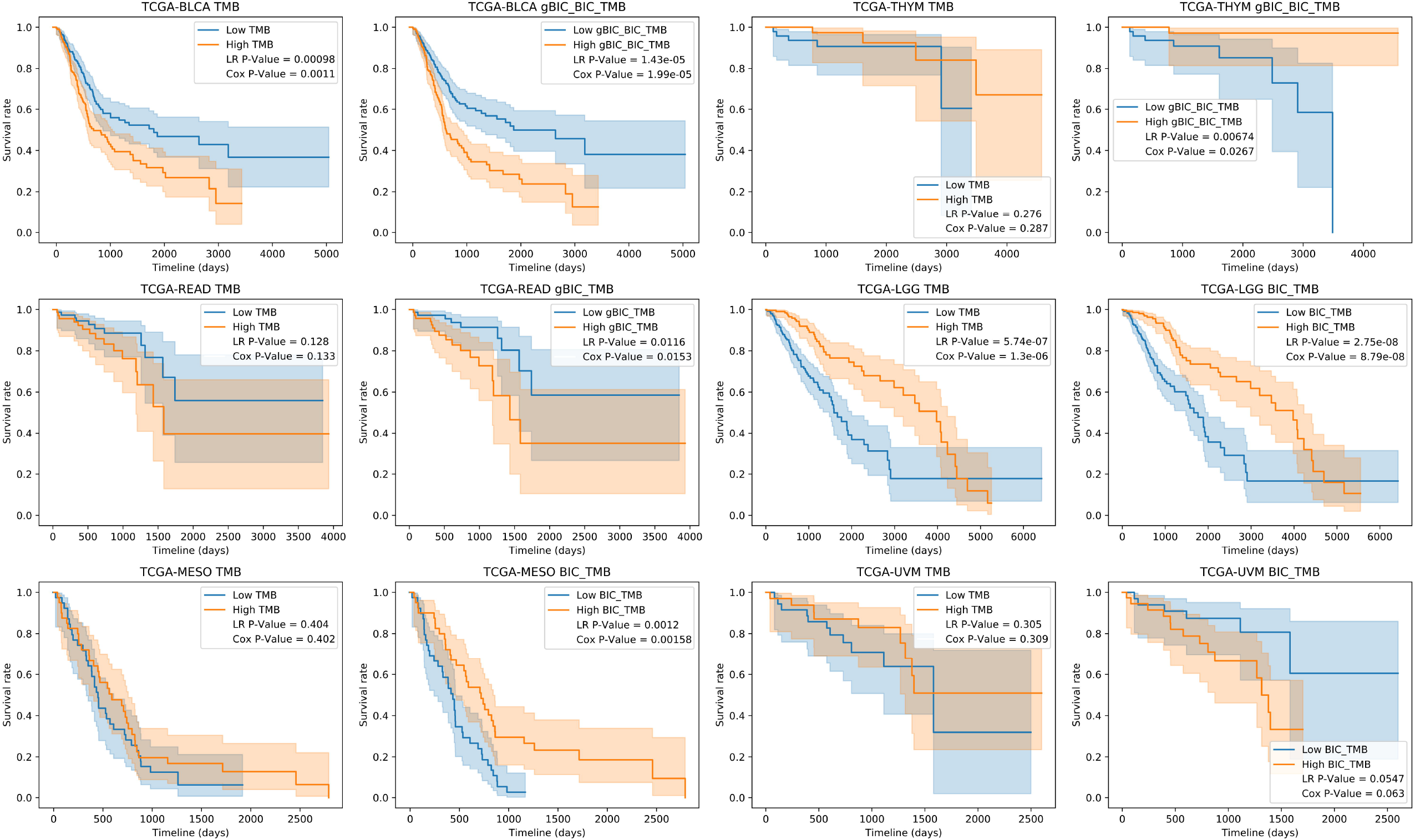
The survival curve of a) BLCA, b) THYM, c) READ, d) LGG, e) MESO and f) UVM grouping by original tumor mutation burden (TMB) and BIC derived TMB. BLCA, Bladder Urothelial Carcinoma; THYM, Thymoma; READ, Rectum adenocarcinoma; LGG, Brain Lower Grade Glioma; MESO, Mesothelioma.

### BIC is able to predict the transmissibility of SARS-CoV-2 strains

COVID-19 is an epidemic disease caused by a severe acute respiratory syndrome coronavirus 2 (SARS-CoV-2), which is a single-stranded RNA virus of the genus Betacoronavirus^29^. Recently, it has been proven that the gene importance (GIC) score can successfully predict the death rate of COVID-19^30^. This suggest that single base mutations in SARS-CoV-2 genome could shape its features such as transmissibility. To confirm this, we explored the correlation between transmissibility of different SARS-CoV-2 lineages and their corresponding BIC score. Doubling Time is the projected time (in months) the virus would take to double the number of samples for a lineage. The results indicate that average BIC score has a significantly positive correlation with the SARS-CoV-2 doubling time (Rho=0.322, p-value=6.68e-9, Figure 5), meaning that SARS-CoV-2 with higher average BIC score on its mutation sites would have greater transmissibility. As we have demonstrated that the BIC score can measure the pathogenicity, which further suggests that SARS-CoV-2 strains with higher average BIC scores would cause higher mortality and lower transmissibility.

**Figure 5.**
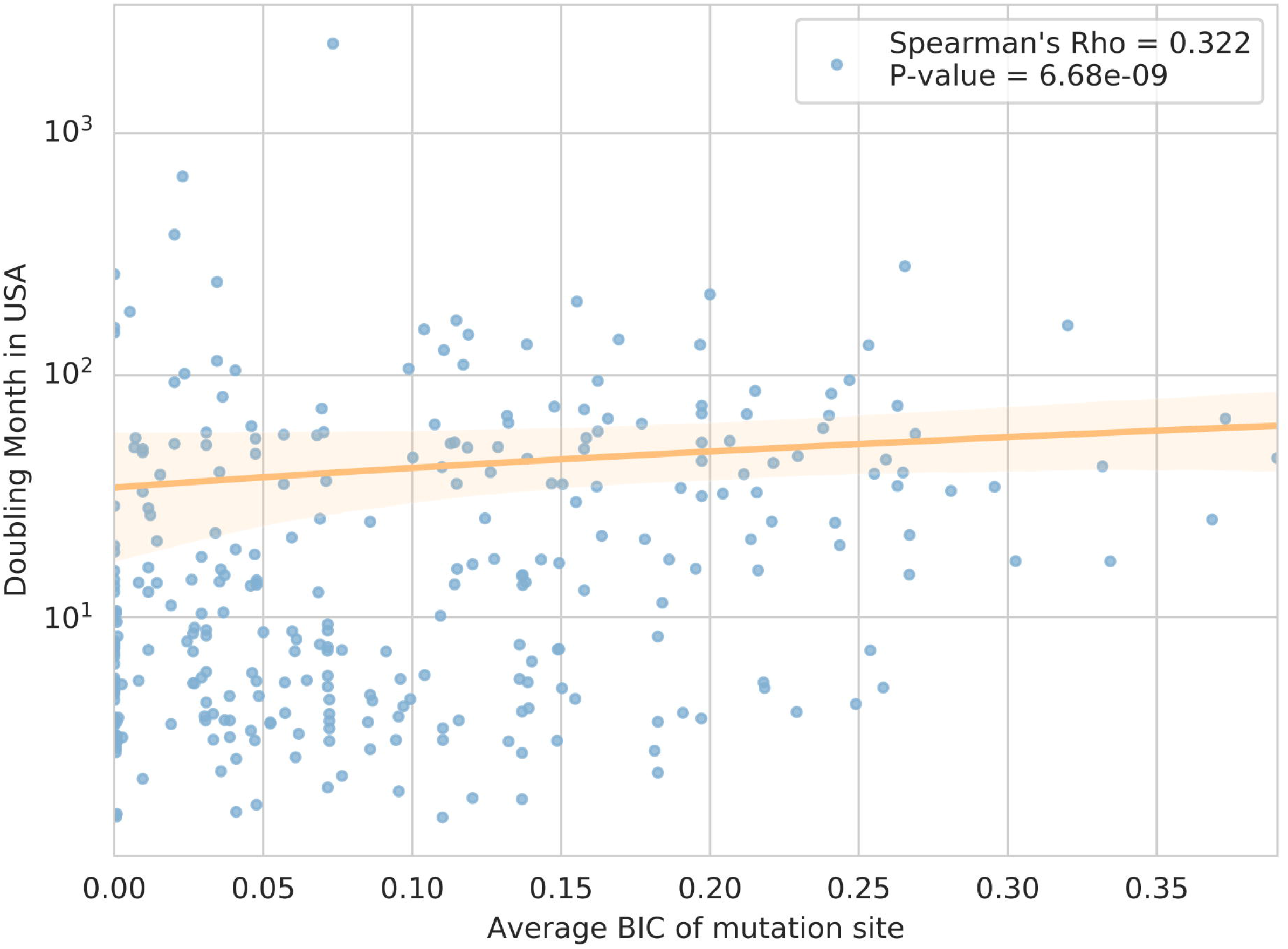
The correlation between average BIC score and SARS-CoV-2 transmissibility. The p-value was calculated by Spearman’s correlation analysis.

### Web server construction

Finally, we have developed a user-friendly online tool that is freely accessible on our website (http://www.cuilab.cn/bic/). This tool allows users to input one mRNA or lncRNA sequence and obtain the BIC score for each nucleotide in the sequence. The results can be downloaded from the result page. Furthermore, we also developed a standalone version of BIC, which is written in Python 3 and can be downloaded from our website.

## DISCUSSION AND CONCLUSION

As the fundamental unit of a gene and its transcripts, nucleotides have enormous impacts on molecular function and evolution, and thus on phenotypes and diseases. Given that different nucleotides on one gene often exhibit diverse levels of effects, it is quite crucial to comprehensively and quantitatively measure the importance of each base on a gene transcript. Although some tools have been developed for evaluating the pathogenicity of missense variants, one class of specific bases representing only a very small fraction of the bases in mRNAs, tools for all bases on mRNAs and lncRNAs are still not available. In this paper, we proposed an algorithm, base importance calculator (BIC), to comprehensively and quantitatively measure the importance score of each single base on human mRNAs and lncRNAs. We further found that BIC can predict the pathogenicity of mutations on at both the base- and the gene-levels. Moreover, we revealed that BIC enhanced the prognosis prediction of a number of cancers. Finally, we found that gBIC has a significantly positive correlation with the transmissibility of SARS-CoV-2 strains. The above results confirmed the effectiveness of the BIC algorithm. Besides its effectiveness, BIC is quite applicative and easy to use as it solely relies on the sequence of mRNAs or lncRNAs.

In addition, besides the proposed three scientific tasks, BIC would contribute to a number of sequence-based bioinformatics tools and analysis. For instance, the BIC score may be useful in identifying pathogenic mutations during GWAS analysis. Recently, a weighted version of the enrichment analysis tool (WEAT) was proposed, taking gene importance scores into account in enrichment analysis^31^. As a gene-level pathogenicity metric, gBIC can serve as a weight for genes to enrich pathways that are more related to diseases. Another example could be metrics in molecular evolution. Tools such as dNdS is dependent on a hypothesis that synonymous mutation is neutral. In the framework of single base importance, the hypothesis could be not such true ^32,33^ and should be improved as well.

Overall, this study has developed BIC algorithm to measure the importance of each single base on a human mRNA or lncRNA sequence. We highlight the potential usefulness of BIC algorithm through three different tasks. In the future, BIC should be confirmed comprehensively by more tasks and could be used to improve a number of sequence-based bioinformatics studies.

## FUNDING

This work was supported by the National Science Foundation of China (Nos. 62072154 and 81921001).

### Author contributions

QC presented the original idea. CC and RF designed the study. RF performed the study. XJ processed the TCGA mutation data. RF, QC, CC, and JL wrote or edited the manuscript.

## Figure Legends

**Supplementary Figure 1.**
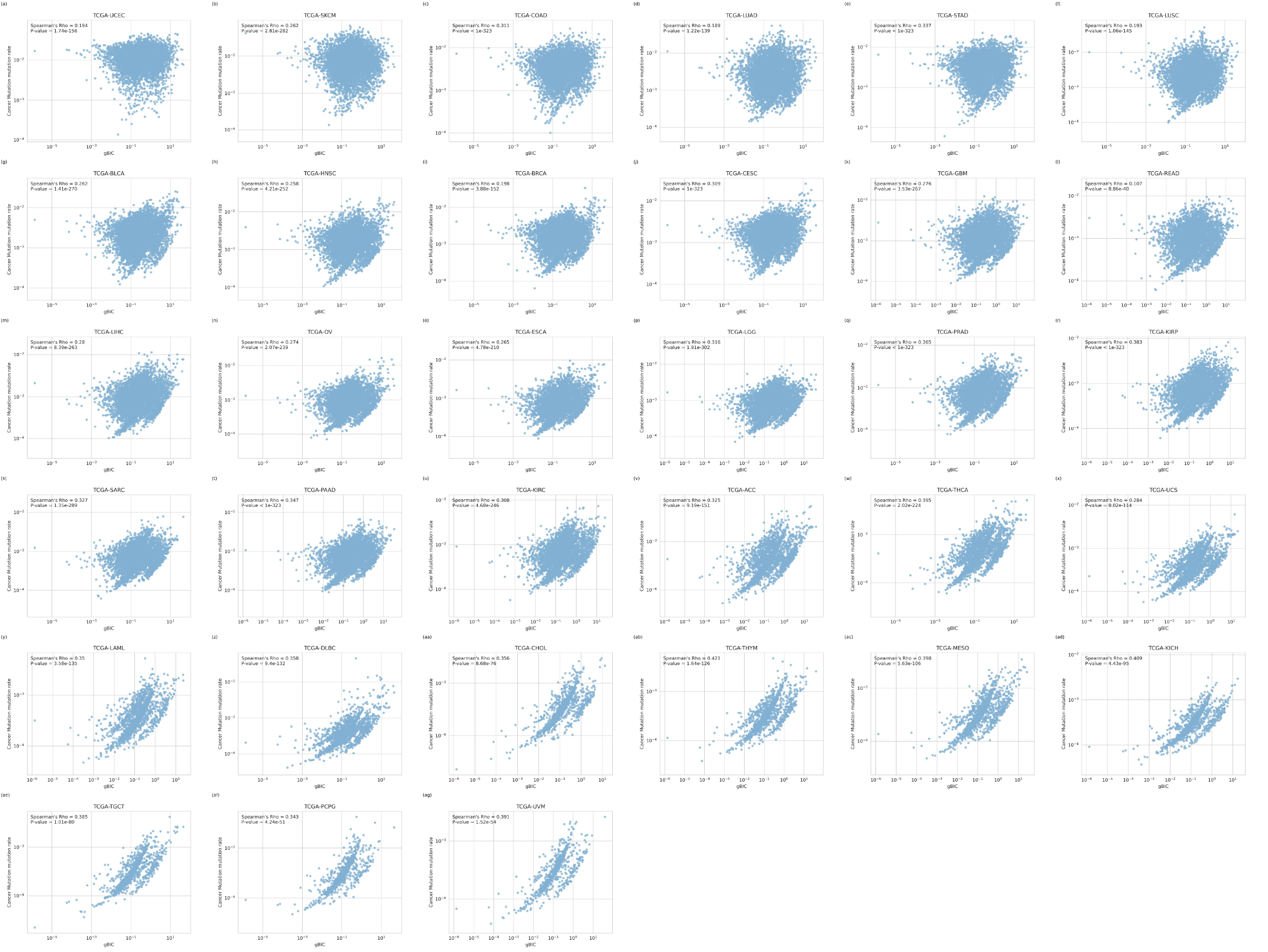
The correlation between the gene level BIC (gBIC) score and the somatic mutation rate of all cancers in TCGA. ACC, Adrenocortical carcinoma; BLCA, Bladder Urothelial Carcinoma; BRCA, Breast invasive carcinoma; CESC, Cervical squamous cell carcinoma and endocervical adenocarcinoma; CHOL, Cholangiocarcinoma; COAD, Colon adenocarcinoma; ESCA, Esophageal carcinoma; GBM, Glioblastoma multiforme; HNSC, Head and Neck squamous cell carcinoma; KICH, Kidney Chromophobe; KIRC, Kidney renal clear cell carcinoma; KIRP, Kidney renal papillary cell carcinoma; LAML, Acute Myeloid Leukemia; LGG, Brain Lower Grade Glioma; LIHC, Liver hepatocellular carcinoma; LUAD, Lung adenocarcinoma; LUSC, Lung squamous cell carcinoma; MESO, Mesothelioma; OV, Ovarian serous cystadenocarcinoma; PAAD, Pancreatic adenocarcinoma; PCPG, Pheochromocytoma and Paraganglioma; PRAD, Prostate adenocarcinoma; READ, Rectum adenocarcinoma; SARC, Sarcoma; SKCM, Skin Cutaneous Melanoma; STAD, Stomach adenocarcinoma; TGCT, Testicular Germ Cell Tumors; THCA, Thyroid carcinoma; THYM, Thymoma; UCEC, Uterine Corpus Endometrial Carcinoma; UCS, Uterine Carcinosarcoma; UVM, Uveal Melanoma; DLBC, Lymphoid Neoplasm Diffuse Large B-cell Lymphoma.

**Supplementary Figure 2.**
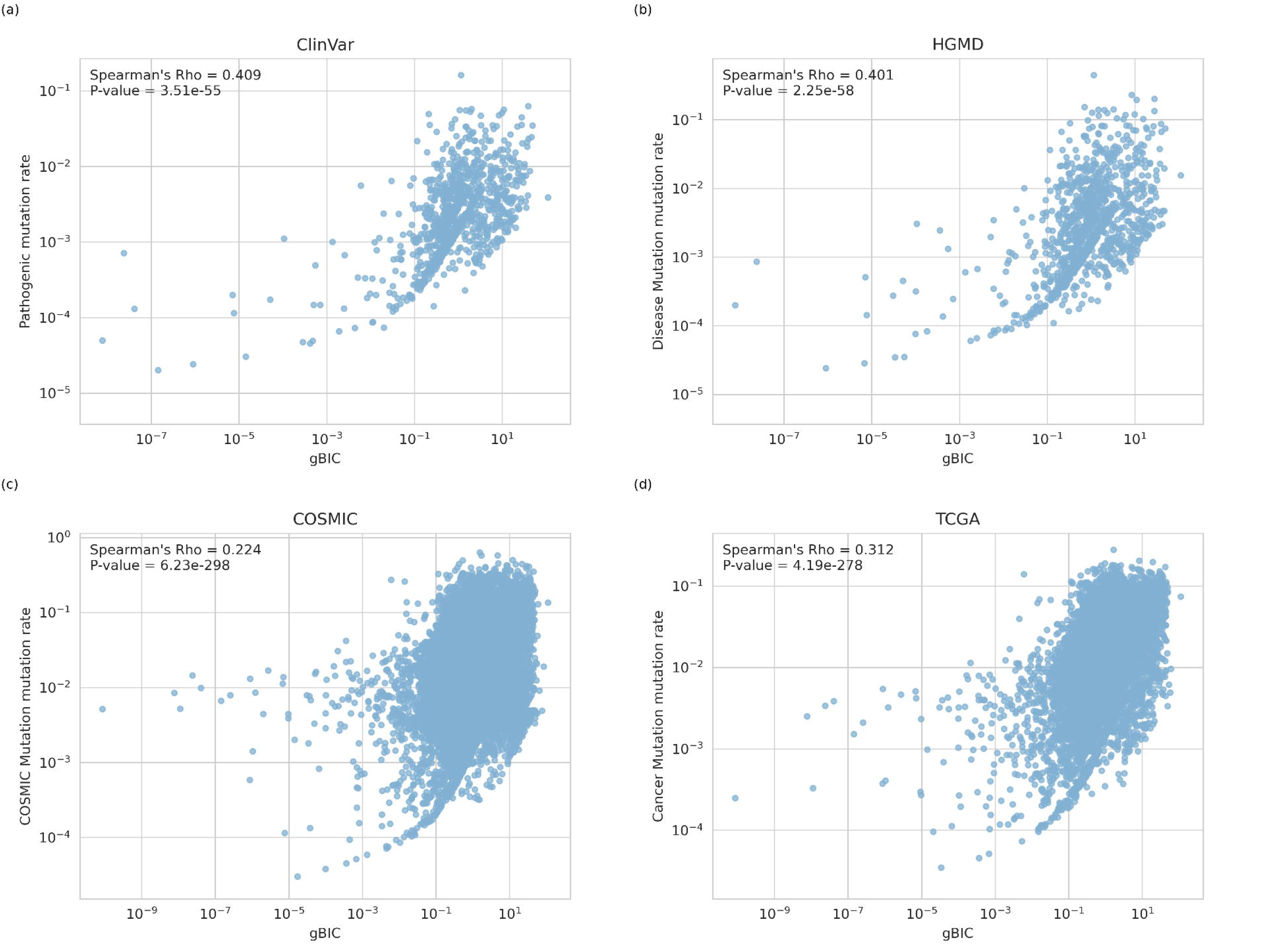
The correlation between gene-level BIC (gBIC) score and the pathological mutation rate for lncRNAs in databases of a) ClinVar, b) HGMD, c) COSMIC, and d) TCGA. The p-value was calculated by Spearman’s correlation analysis.

**Supplementary Figure 3.**
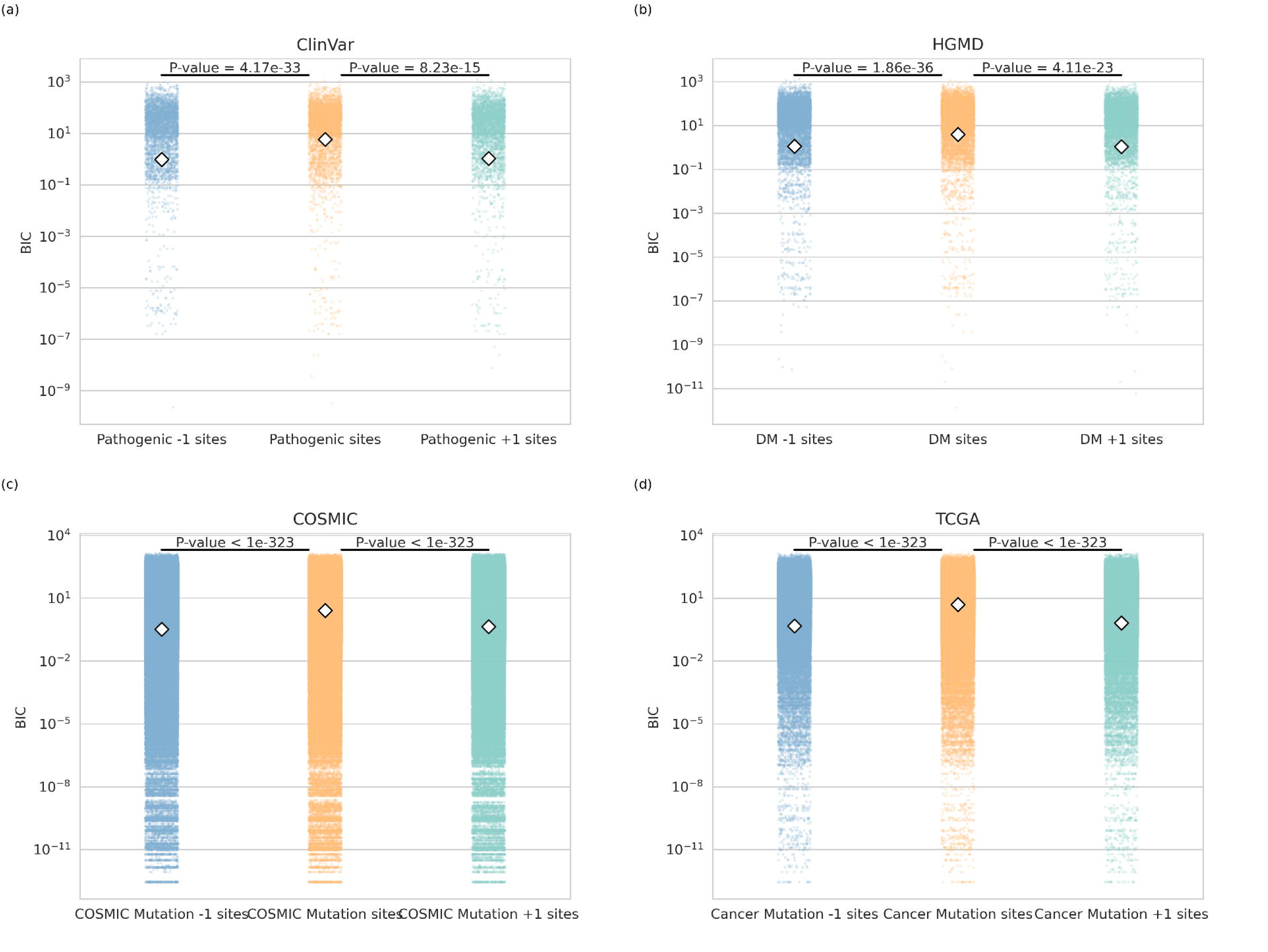
Comparison of the BIC scores of the bases hosting pathological mutation sites (yellow) and their upstream neighbor single bases (blue) and downstream neighbor single bases (green) for lncRNAs in databases of a) ClinVar, b) HGMD, c) COSMIC, and d) TCGA. The p-value was calculated by Mann-Whitney U rank test.

**Supplementary Figure 4.**
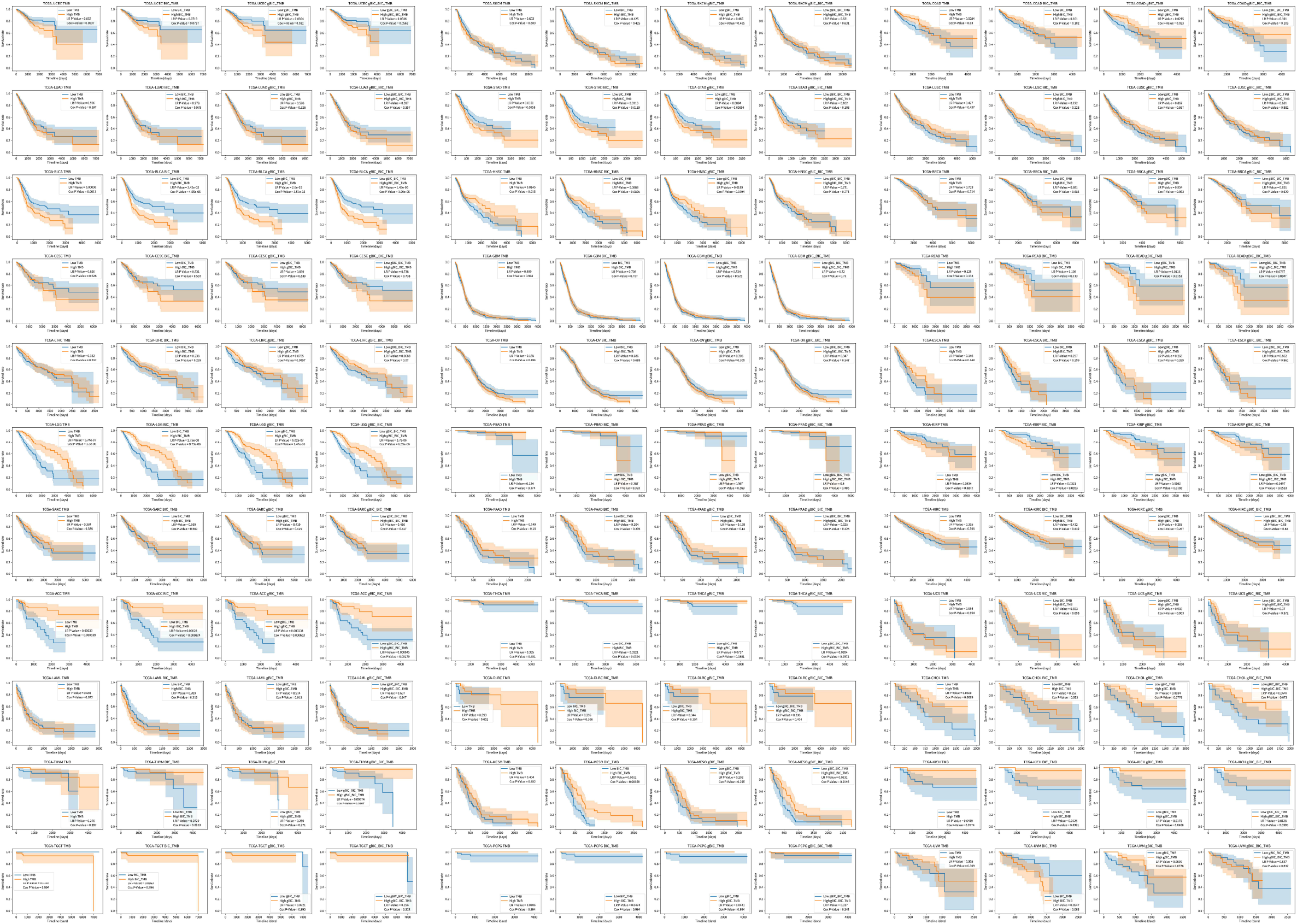
The survival curve of all TCGA cancers grouping by original tumor mutation burden (TMB), gBIC weighted TMB, BIC weighted TMB and both gBIC and BIC weighted TMB. ACC, Adrenocortical carcinoma; BLCA, Bladder Urothelial Carcinoma; BRCA, Breast invasive carcinoma; CESC, Cervical squamous cell carcinoma and endocervical adenocarcinoma; CHOL, Cholangiocarcinoma; COAD, Colon adenocarcinoma; ESCA, Esophageal carcinoma; GBM, Glioblastoma multiforme; HNSC, Head and Neck squamous cell carcinoma; KICH, Kidney Chromophobe; KIRC, Kidney renal clear cell carcinoma; KIRP, Kidney renal papillary cell carcinoma; LAML, Acute Myeloid Leukemia; LGG, Brain Lower Grade Glioma; LIHC, Liver hepatocellular carcinoma; LUAD, Lung adenocarcinoma; LUSC, Lung squamous cell carcinoma; MESO, Mesothelioma; OV, Ovarian serous cystadenocarcinoma; PAAD, Pancreatic adenocarcinoma; PCPG, Pheochromocytoma and Paraganglioma; PRAD, Prostate adenocarcinoma; READ, Rectum adenocarcinoma; SARC, Sarcoma; SKCM, Skin Cutaneous Melanoma; STAD, Stomach adenocarcinoma; TGCT, Testicular Germ Cell Tumors; THCA, Thyroid carcinoma; THYM, Thymoma; UCEC, Uterine Corpus Endometrial Carcinoma; UCS, Uterine Carcinosarcoma; UVM, Uveal Melanoma; DLBC, Lymphoid Neoplasm Diffuse Large B-cell Lymphoma.

**Supplementary Figure 5.**
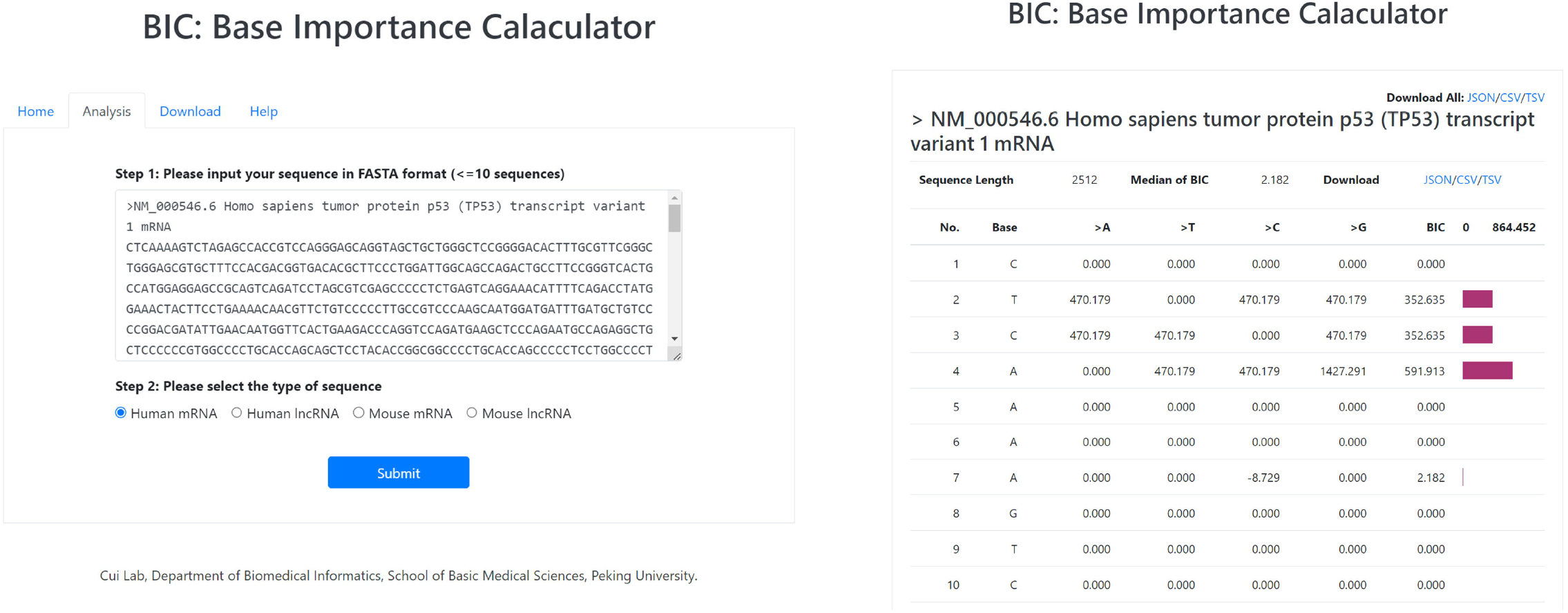
The web interface of BIC online tool.

